# Compression-based Network Interpretability Schemes

**DOI:** 10.1101/2020.10.27.358226

**Authors:** Jonathan Warrell, Hussein Mohsen, Mark Gerstein

## Abstract

Deep learning methods have achieved state-of-the-art performance in many domains of artificial intelligence, but are typically hard to interpret. Network interpretation is important for multiple reasons, including knowledge discovery, hypothesis generation, fairness and establishing trust. Model transformations provide a general approach to interpreting a trained network post-hoc: the network is approximated by a model, which is typically compressed, whose structure can be more easily interpreted in some way (we call such approaches *interpretability schemes*). However, the relationship between compression and interpretation has not been fully explored: How much should a network be compressed for optimal extraction of interpretable information? Should compression be combined with other criteria when selecting model transformations? We investigate these issues using two different compression-based schemes, which aim to extract orthogonal kinds of information, pertaining to feature and data instance-based groupings respectively. The first (*rank projection trees*) uses a structured sparsification method such that nested groups of features can be extracted having potential joint interactions. The second (*cascaded network decomposition*) splits a network into a cascade of simpler networks, allowing groups of training instances with similar characteristics to be extracted at each stage of the cascade. We use predictive tasks in cancer and psychiatric genomics to assess the ability of these approaches to extract informative feature and data-point groupings from trained networks. We show that the generalization error of a network provides an indicator of the quality of the information extracted; further we derive PAC-Bayes generalization bounds for both schemes, which we show can be used as proxy indicators, and can thus provide a criterion for selecting the optimal compression. Finally, we show that the PAC-Bayes framework can be naturally modified to incorporate additional criteria alongside compression, such as prior knowledge based on previous models, which can enhance interpretable model selection.

## 1. Introduction

Numerous methods have been proposed to address the problem of model interpretability in the context of deep learning [1]. These include embedding structure relevant to a problem in the network architecture [2-4], modifying the training objective to include a cost relevant to interpretability [5,6], and post-hoc extraction of features of feature-groups which have been prioritized by the network [7-12] Moreover, interpretations can aim to explain aspects of the model globally [12], or with respect to the predictions of individual instances [13,14]. Here, we will focus on methods where the aim is to perform post-hoc interpretation on a model globally, by applying a compression-based model transformation which allows domain-relevant information to be extracted relevant to how the model is making predictions. Although compression techniques may be used solely to enhance comprehension [13,14], our interest is in using compression both for this purpose and as a means of implementing a *minimum description length* (MDL) prior to identify generalizable structure [15]. As we have argued elsewhere [16], interpreting a network may, in general, involve making explicit any aspects of network structure that implicitly relate to domain-relevant phenomena (implicit semantics). A principled way of finding such aspects is to identify structure that is expected to generalize by using MDL as a generalized form of Occam’s razor.

Network compression has been studied previously from multiple different perspectives. The most direct motivation is to create an efficient representation of a larger network suitable for limited memory and/or running time applications [17]. Another area of interest has been the application of compression methods to provide bounds on the generalization error of a network [15,18]. Particularly, tight bounds have been shown to be possible in this setting using a PAC-Bayes framework [15]. A related application of compression techniques is the control of generalization through regularization [19,20]. In the field of network interpretability, model reduction techniques have been used to find a model in a simpler class, which may be used to approximate a more complex network [13,14,21]. The simpler model may be chosen from a class such as linear models, which are more easily interpreted than the original model. Models for interpretability formulated this way leave many aspects open, such as the degree and type of compression, to be specified by the user, in response to a tailored definition of interpretability for an individual problem. However, as outlined above, we are interested in a general definition of interpretability, capable of identifying domain-relevant structure in the network’s architecture, without restriction to a particular target model class. Further, in this context a number of additional questions arise, such as how much a network should be compressed for optimal extraction of interpretable information, and whether compression be combined with other criteria when selecting model transformations. We propose that techniques for network compression and generalization analysis such as above may be combined to offer a principled solution to these issues. Indeed, the relationship between generalization and interpretability has not been characterized in previous work. In order to investigate these issues in a general setting, we introduce the notion of an *interpretability scheme*, consisting of a) a method of model transformation/compression applied to a pretrained network, b) a method for extracting interpretable information from the transformed model, and c) a (PAC-Bayes based) method for scoring transformed models for interpretable model selection (see Fig. 1A; we note that the pretrained network may or may not embed aspects of prior domain knowledge).

**Figure 1.**
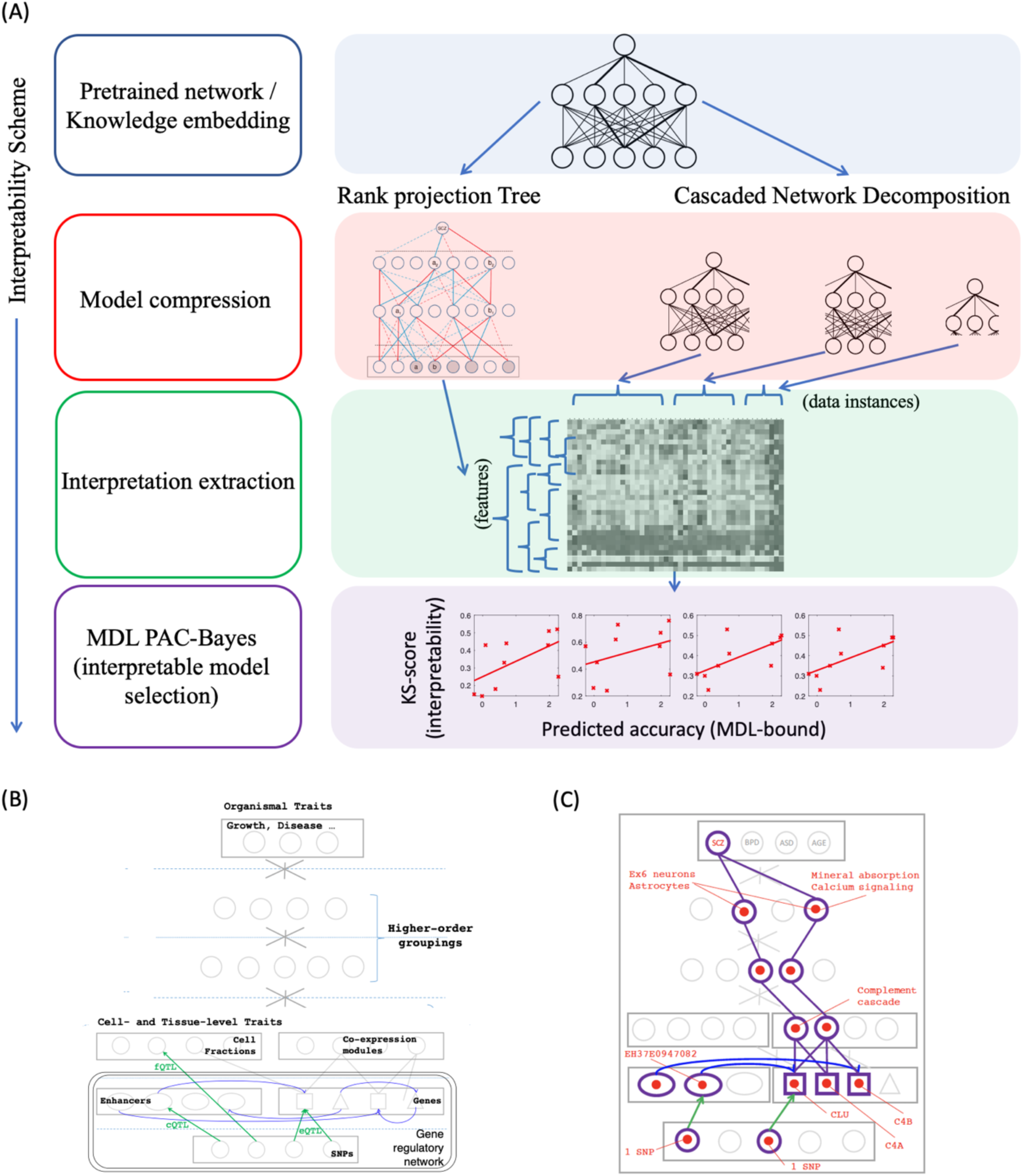
Compression-based interpretability schemes. (A) Illustrates general framework, and orthogonal Rank Projection Tree (RPT) and Cascaded Network Decomposition (CND) interpretability schemes investigated in paper. (B) Shows example of DSPN network used in PsychENCODE [2], in which base layers are trained as Boltzmann machine with connectivity used to embed prior biological structure, upper layers as a feed-forward network, and RPT scheme is used to infer higher-order groupings of genes and other biological entities. (C) Shows example interpretation, in which RPT is used to infer groupings, modules, genes, enhancers and SNPs associated with Schizophrenia. See [2] for further biological details.

We explore two different interpretability schemes in the following to show the generality of our approach. The first (*rank projection trees*, RPTs) uses a structured sparsification method that nested groups of features can be extracted having potential joint interactions. Previous interpretation methods for prioritizing features include gradient-based schemes [7,8], gradient+input schemes [9], perturbation schemes [10,11], and difference scores with respect to a reference [12]. All such methods involve scoring individual nodes at a layer of interest, while we may be interested in picking out a collection of subsets of nodes at multiple scales from a given layer which are ‘important’ to the network. Such a situation often arises in biological applications where we are interested in epistatic effects between groups of genes in determining a trait of interest. For instance, the framework of weighted gene coexpression network analysis (WGCNA, see [22,23]) is widely used to group together genes sharing common coexpression patterns. However, WGCNA does not use trait-relevant information in defining modules.

Additionally, methods based on Shapley features [14] pick out feature groupings, but do not explicitly derive these from the network structure. Our RPT approach can be used to select collections of trait-relevant subsets of interacting genes (and other elements) by providing a multiscale interpretation of a supervised deep neural network. Further, the framework is agnostic as to the underlying ranking function which is used to build the tree, allowing any of the above feature prioritization methods to be used to provide a scoring function.

The second interpretability scheme we investigate (*cascaded network decomposition*, CND) is orthogonal to the first, in focusing on groups of data instances as opposed to feature groupings. Here, we identify groups of training (or testing) instances with similar properties through compressing a pretrained network in a series of stages to form a cascade, where each stage accepts and rejects subsets of the training data, and identifies a subnetwork of the original model that focusses on classifying these instances (for instance, for a face classifier trained to recognize gender, the subnetworks may focus on different ages or ethnicities). The instances accepted at each stage form a group with a similar level of ‘difficulty’. The approach may be viewed as a form of boosting [24], and has similarities with Deep Boosting [25] and Deep Cascade [26] approaches, which use *de novo* trained deep networks as the base and/or internal node classifiers in a cascade model. Both approaches [25,26] also derive Radamacher complexity bounds on the generalization error for the full cascade. In contrast, we use deep networks extracted from a pre-existing trained model as the base classifiers, and internal node classifiers which reflect groupings of data-points implicit in the trained model. Further, we analyze generalization performance using MDL PAC-Bayes bounds, as discussed.

We use a common framework for analyzing the generalization performance of both schemes discussed above, which we show can be used for interpretable model selection in the experimentation. For each scheme, we derive an MDL PAC-Bayes bound by adapting the approach of [15] to use the scheme-specific compression schemes we introduce. Further, as noted above, we are interested in whether compression can be combined with other criteria when selecting model transformations. Particularly, previous PAC-Bayes approaches have shown that data-dependent priors may be used to provide more informative bounds, allowing prior knowledge of a domain to inform model selection [27,28]. We thus derive a novel bound which combines both data-dependent and MDL components, hence trading off compression and domain knowledge, and show how this can be applied to the schemes we introduce.

For our empirical investigation, we use the schemes introduced to interpret networks trained on cancer and psychiatric genomics classification tasks, using genomics data from PCAWG [29] and PsychENCODE [30,2] consortia. We show that the RPT scheme is able to pick out biologically important genes and gene sets in both domains, using networks trained to predict epistatic interactions of germline and somatic mutations in cancer, and risk for Schizophrenia, respectively. In doing so, we extend the results of [2], who show that the RPT can be used to find groupings of higher-order groupings genes and elements in a network (DSPN) specifically designed to embed prior biological network structure to aid interpretability (Fig. 1B-C). Further, we show that the CND scheme is able to extract groupings of patients corresponding to covariates (gender, age, ethnicity), medical (medication usage) and disease-subtype categories for Schizophrenia, Bipolar and Autism subjects. Finally, we show that network generalization is informative for interpretation quality using both schemes, and that the MDL PAC-Bayes bounds we introduce can be used as proxy scores for interpretable model selection, hence supporting the general applicability of compression-based interpretability schemes.

In summary, we introduce two orthogonal approaches to compression-based interpretation (RPT and CND), in addition to a novel data-dependent compression bound using a PAC-Bayes approach. Further, our experimentation demonstrates the feasibility of compression-based interpretability schemes using generalization bounds. We begin in Sec. 2 by outlining our interpretability schemes and associated PAC-Bayes generalization bounds, before describing our empirical investigation using cancer and psychiatric genomics classification tasks in Sec. 3. Sec. 4 then concludes with a discussion.

## 2. Compression-based Interpretability

As introduced in Sec. 1, an *interpretability scheme*, consists of a) a method of model transformation/compression, b) a method for extracting interpretable information from the transformed model, and c) a (PAC-Bayes based) method for scoring transformed models for interpretable model selection. Here, we introduce two orthogonal methods for compression-based interpretability, describing the associated compression and extraction schemes for each in Secs. 2.1-2, and common methods for PAC-Bayes based model selection in Sec. 2.3.

### 2.1 Rank Projection Trees

We assume we have a neural network *N* with layers *l* = 0… *L*, where layers 0 and *L* are the output and input layers, respectively (note that the inputs may be an interpretable layer in a larger network, such as the DSPN, where they are an embedded layer incorporating biological network structure, see Fig. 1B-C). Each layer *l* has an associated set *I*_*l*_ = {1… *N*_*l*_} indexing the nodes of that layer; hence *n*_*l*.*i*_ is the *i*’th node at layer *l*. For convenience, we assume there is one output node, *n*_0_,_1_. We write the weight matrix between layers *l*_1_ and *l*_2_ = *l*_1_ − 1 as 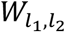 and the biases vector at layer *l* as *β*_*l*_. A *rank projection tree* (RPT, see Fig. 2) over a given network is fully determined by specifying (a) a half branching factor 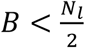,∀,*l* and (b) a ranking function *r*_*i*_,_*l*_,_*m*_(*j*), where *l < m* ≤ *L* are layer indices, *i* and *j* are node indices on layers *l* and *m* respectively, and the function returns an integer specifying the position of node *j* in an ordering of the nodes at layer *m* according to their ‘score’ with respect to node *i* and layer *l*. Semantically, we expect that increased activation of node *j* towards the top of the ranking will lead to increased activation of node *i*, while increased activation of nodes towards the bottom will lead to decreased activation of *i*; hence, any score function of the kind described above (such as the gradient) may be used.

**Figure 2.**
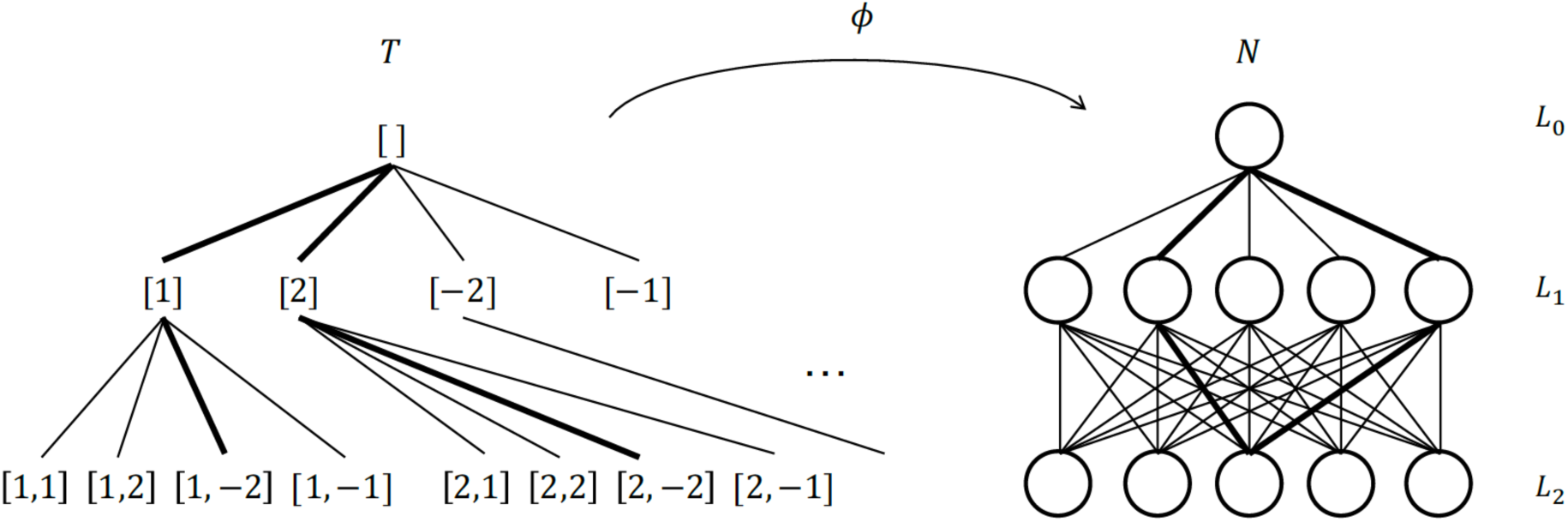
Rank projection trees. The rank projection tree (left, *T*) is mapped onto a trained neural network (right, *N*) via the mapping *ϕ* which depends on an arbitrary ranking function *r*. The image of *T* under *ϕ* is used to prioritize inputs and sets of inputs in *N* in an output dependent fashion.

The nodes of the rank projection tree *T* are lists of ‘branching indices’ of the form [], [*b*_1_], [*b*_1_, *b*_2_],…[*b*_1_,…, *b*_*L*_], where *b*_*l*_ ∈ {1,…, *B*} ∪ {−1,…, −*B*}, ∀ *l*. The node [] is the root of the tree, and the parent function is defined as *Pa*([*b*_1_,…, *b*_*l*−1_, *b*_*l*_]) = [*b*_1_,…, *b*_*l*−1_]. A node *t* of *T*, where *t* is a list of length *l*, is then associated with a node in layer *l* of the neural network via a function *ϕ* defined recursively as follows:

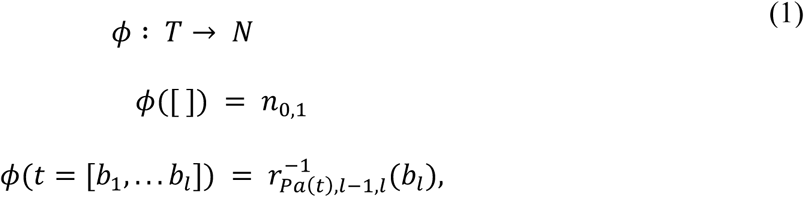

where 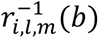 is a ‘quasi-inverse’ of the ranking function, which returns the node *n*_*m*_,_*j*_ for which *r*_*i*_,_*l*_,_*m*_(*j*) = *b* if *b* > 0, and *n*_*m*_,_*j*_ for which *r*_*i*_,_*l*_,_*m*_(*j*) = *N*_*m*_ + *b* + 1 if *b <* 0. For node *t* in *T* then, which maps to *ϕ*(*t*) = *n* at layer *l*, the mappings of the children of *t* are set by first ranking layer *l* + 1 of the neural net with respect to *n*, and assigning the top *B* and bottom *B* nodes of this ranking to the children of *t*, hence *projecting* the full ranking onto a reduced ranking across the 2*B* children.

Each node *t* in *T* may be associated with positive and negative subsets, 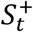 and 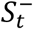, at a reference layer, which we take to be the input layer *L*. These are defined as:

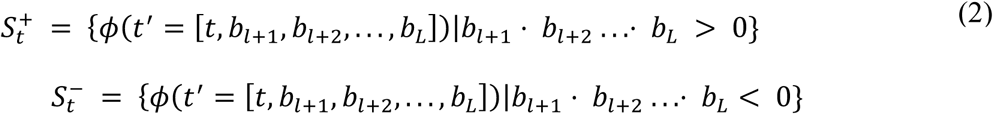

Hence, 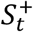 contains all those nodes mapped to by descendants of *t* at layer *L* along paths where the product of the branching indices below *l* is positive, and 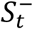 is defined similarly, but where the product of the branching indices is negative. A collection of ‘prioritized’ subsets at multiple levels is thus formed by applying Eq. 2 as *t* runs across *T*. We note that, since multiple nodes in *T* may map to the same node in *N*, sets at the same layer may overlap, including positive and negative sets associated with the same node in *T*. Finally, we may define a prioritization function *π* (or ‘salience map’) of the nodes at the reference layer in *N, π*(*n*) = *f*(*ϕ*^−1^(*n*)),

#### Algorithm 1. Cascaded Network Decomposition (CND)

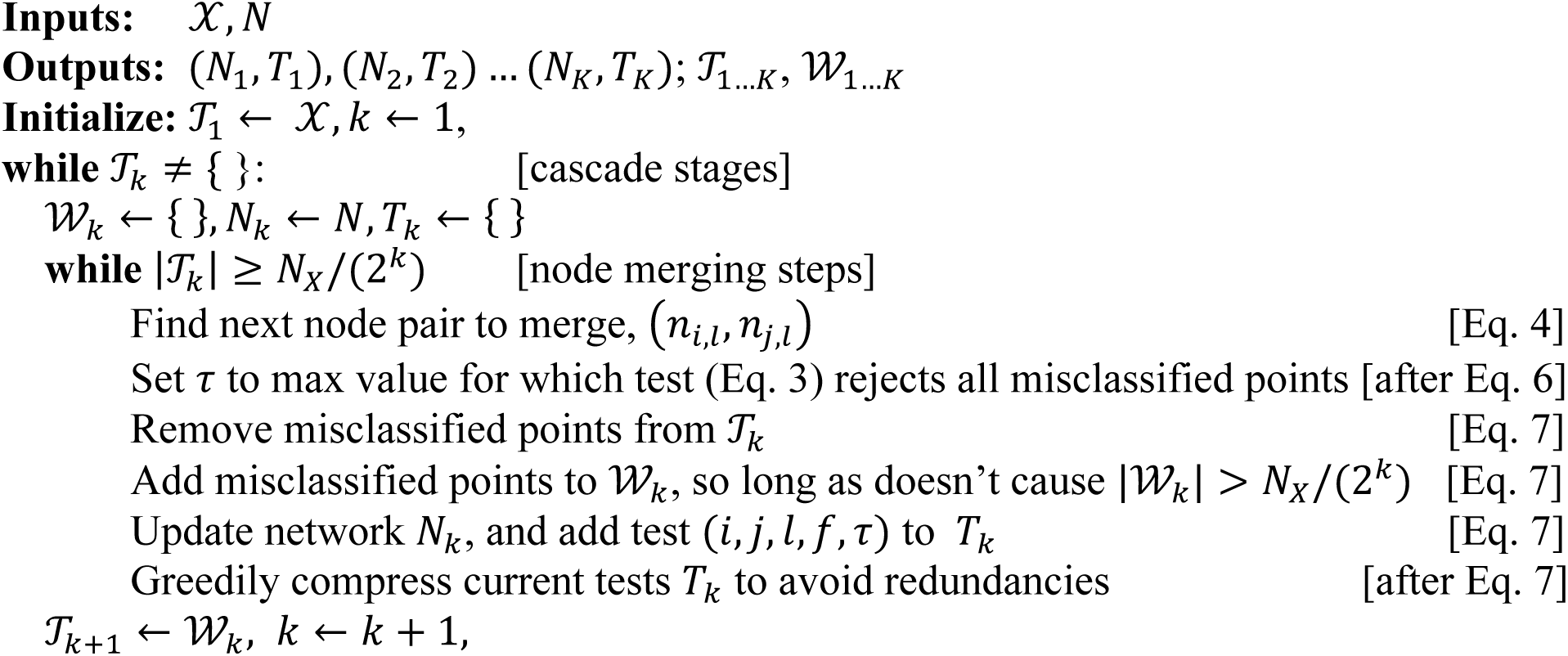

where *ϕ*^−1^(*n*) = {*t*_1_, *t*_2_…} is the pre-image of *n* under *ϕ*, and *f* may be chosen from a number of possibilities (see Sec. 3.1).

### 2.2 Cascaded Network Decomposition

We again assume we have a trained neural network *N* with layers *l* = 0 … *L*, where layers 0 and *L* are the output and input layers, respectively, and *n*_*l*.*i*_ is the *i*’th node at layer *l*. For the cascaded network decomposition (CND) algorithm, we also assume we have access to the training data, 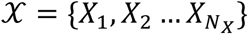. The goal of CND is to provide a cascade of networks, *N*_1_, *N*_2_ … *N*_k_, along with a set of rejection tests at each stage of the form 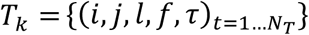. A data-point is fed through each stage of the cascade sequentially (see Fig. 3A); at stage *k*, for each test the following Boolean test is evaluated:

**Figure 3.**
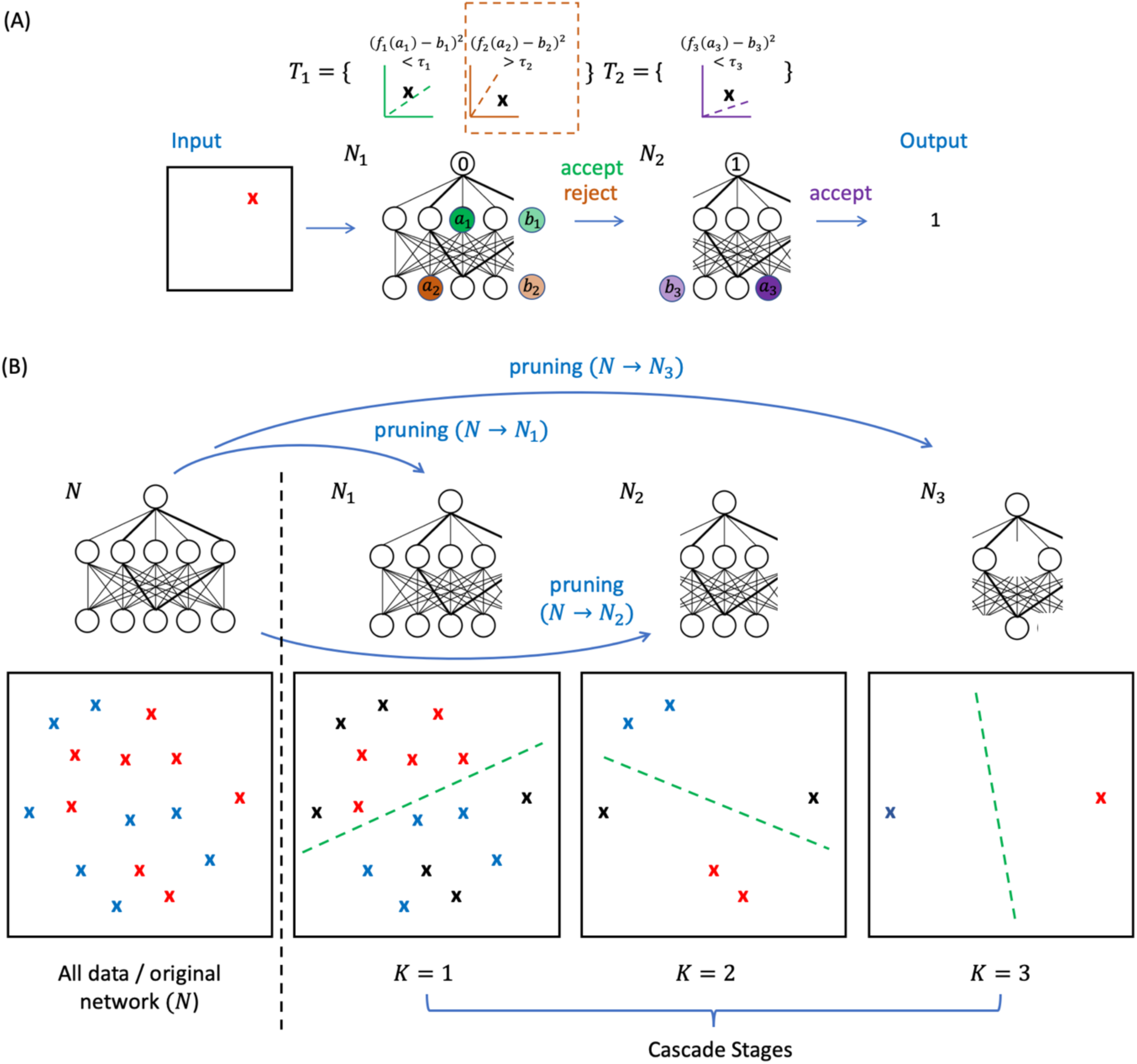
Cascaded Network Decomposition. (A) illustrates the cascade classifier applied to a datapoint. At stage 1 of the cascade, the datapoint is fed through network *N*_1_, giving output 0; further, two tests belonging to *T*_1_ are applied, each comparing a merged node pair in *N*_1_ (where the second node (*b*) is a ‘ghost’ node with no outputs; note that for test (*i, j, l, f, τ*)_t_, *a*_t_ = *a*_*i*_,_*l*_,_k_ and *b*_t_ = *a*_1_,_*l*_,_k_). Since the second test signals rejection (*f*_2_(*a*_2_) − *b*_2_)^2^ > *τ*, see dotted box), the datapoint is passed to stage 2 of the cascade. Here, the single test in *T*_2_ is accepted, and so the cascade outputs the result of *N*_2_ and predicts class 1. (B) illustrates CND training algorithm. The algorithm begins with a pre-trained neural network (*N*) which perfectly classifies the full dataset (left). The cascade is then built in a series of stages (*K*=1,2,3), where each stage compresses the original network to classify at least half of the data points from the previous stage. The data points which are misclassified at each stage are shown in black, and these are sent on to the next stage of the cascade. The output is a set of compressed networks, one for each cascade stage (*N*_1_, *N*_2_, *N*_9_), and tests for identifying which data points to send to the next stage (*T*_1_, *T*_2_).

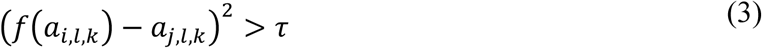

where *a*_*i*_,_*l*_,_k_ is the activation of the *i*’th node at layer *l* in network *N*_k_ evaluated on the given data-point, and *f* is a *matching function*, defined below. If the test in Eq. 3 is true for any of the tests at stage *k*, the data-point is rejected by that stage, and passed onto the next, otherwise the output of *N*_k_ is returned, and no further stages of the cascade are evaluated.

We now describe the algorithm used to derive the networks *N*_1_, *N*_2_ … *N*_k_ and associated tests for each stage (see Fig. 3B for schematic). At each stage of the algorithm, we keep track of a *working set* of training examples, *W*_k_, the *current training set 𝒯*_k_, in addition to *N*_k_ and *T*_k_. The working set represents the training instances to be passed onto the next cascade stage. At stage *k* = 1, we initialize *𝒯*_1_ = *𝒳*, while for *k* > 1 we set *𝒯*_k_ = *𝒲*_k_^*−1*^. Further, we initialize *𝒲*_k_ ={}, *N*_k_ = *N, T*_k_ = {}. Then, at a given cascade stage *k*, we perform network compression in a series of steps, by searching over all ordered pairs of nodes, (*n*_*i*_,_*l*_, *n*_1_,_*l*_) for arbitrary *l*, to find the pair for which:

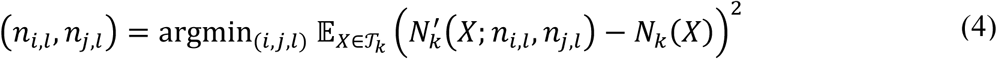

where 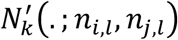 is the network formed from *N*_*k*_ by removing node *n*_*j*_,_*l*_, and transforming the weights and biases:

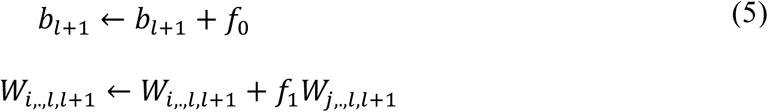

and *f*(*a*) = *f*_0_ + *f*_1_*a* is the linear transform that minimizes the mean squared error:

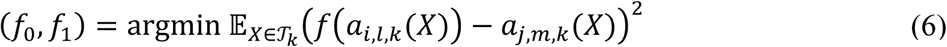

writing *a*(*X*) for an activation evaluated on data-point *X*. For each node merge, a threshold *τ* is set as high as possible, while requiring that for all *X* ∈ *𝒯*_k_, the mean squared error in Eq. 4 is within a specified tolerance *ϵ* whenever the test in Eq. 3 is false (we may set *ϵ* = 0 for binary classification). Then, the following updates are made:

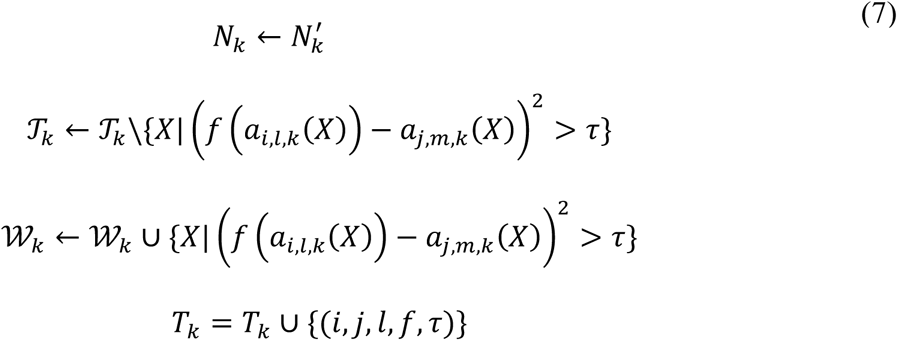

Further, the set *T*_k_ is compressed to find a small set of tests which ‘cover’ the current working set; this may be done greedily, for instance by ordering the tests in descending order of how many points they reject, and successively adding tests only if they reject points not rejected by previous tests. The compression process for stage *k* finishes when it is not possible to merge any more nodes while maintaining the constraint |*𝒲*_k_| ≤ *N*_X_/(2^k^). This ensures that the current training set reduces by at least 1/2 at each stage of the cascade. The final stage *K* of the cascade is reached when *𝒯*_K+1_ = {}. The algorithm ensures that the mean squared error of final cascade classifier will be within *ϵ* of the original network, *N*. Further, we note that when searching over node pairs in Eq. 4, an extra node is included for each bias terms whose activation is always 1, and the input features may further be treated as nodes at level *L* + 1, enabling non-predictive features to be removed in the merging process. Alg. 1 summarizes the algorithm in pseudo-code, which provides the compression method for the CND scheme. The compression scheme results in a partitioning of the training data into the sets of points which are accepted at each stage of the cascade, *𝒯*_*k*_\*𝒲*_*k*_. For the extraction step, these groups of training points may be treated as clusters for further interpretation, either directly, or in a class-specific fashion (hence, for a binary task we extract (*𝒯*_*k*_ \*𝒲*_*k*_) ∩ [*Y* = 0] and (*𝒯*_*k*_ \*𝒲*_*k*_) ∩ [*Y* = 1] for each *k*.

### 2.3 PAC-Bayes Compression Bounds

Finally, we outline the MDL PAC-Bayes bounds that we use to perform interpretable model selection in the experimentation. We use the following basic form of the PAC-Bayes bound, outlined in [31]:

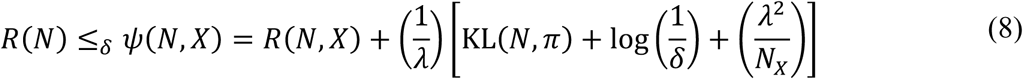

where *R*(*N*_*m*_ is the true risk of network *N, R*(*N, X*) is the empirical risk on the observed sample *X, π* is a prior over networks, and KL(*N, π*) is the KL-divergence of a delta-distribution at *N* with *π*. As introduced in [15], given an encoding scheme, *π* may be formulated as a minimum description length (MDL) prior:

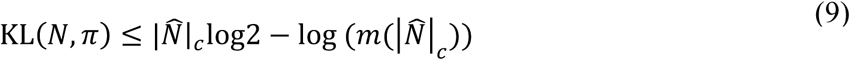

where 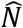 is the code-word for *N* (which may be a lossy code), |. |_*c*_ is the code length, and *m*(. _*m*_ is a prior over code lengths, which for convenience may be taken to be uniform. For RPT and CND schemes, codewords may be generated by taking the compressed representations generated by the schemes in Sec. 2.1 and 2.2 (the sparsified network or sequence of cascade stages plus tests respectively), and subjecting these to further compression via LZW coding to generate a binary code. Eq. 8 can thus be used directly as a generalization bound after substituting Eq. 9 for the KL term.

As noted previously, we are also interested in combining compression with prior information to produce a modified MDL bound. For this purpose, we introduce the following bound (see Appendix A for proof):

#### Theorem 1 (Modified MDL Bound).

Let *π*_*MDL*_ be the MDL prior from [15] (see Th. 4.3, letting *σ*_1_ = *τ*), and *π*_*dep*_ be a data dependent prior, *𝒩*(. ; *N*_0_, *σ*_2_), where *N*_0_ is a pretrained neural network, and *𝒩*(. ; ., *σ*) is a Gaussian with symmetric covariance *σ*. Then, for the weighted prior *π* = *απ*_*MDL*_ + (1 − *α*) *π*_*dep*_ and posterior *ρ* = *𝒩*(. ; *N, σ*_3_) we have:

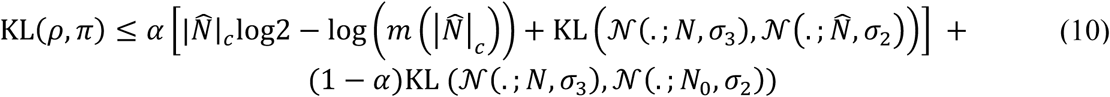

The bound from Eq. 10 can be directly substituted into Eq. 8, after training *N*_0_ on hold-out data following [27,28].

## 3. Empirical Study

In the following, we explore the ability of the *rank projection tree* and *cascaded network decomposition* algorithms defined above to prioritize genes/subsets of genes and groups of subjects/instances relevant to cancer and psychiatric genomics applications. For cancer genomics, we use data from the PanCancer Analysis of Whole Genomes dataset (PCAWG, [29]) to train neural networks (3 hidden layers) to predict the per-tumor co-occurrence of germline and somatic mutations in a given gene (a binary output), using germline variant signatures of known cancer genes alongside a set of gene-level biological features as inputs. For psychiatric genomics, we use data from the PsychENCODE dataset [30,2] to train neural networks (2 hidden layers) to classify control versus Schizophrenia post-mortem subjects after balancing covariates (age, gender, ethnicity, assay), using bulk transcriptomic data from the prefrontal cortex as inputs, either in the form of individual gene expression levels, or average expression across predefined modules of genes (using WGCNA [22,23]). See Appendix A for further details.

### 3.1 Prioritization Functions

We first compare the ability of the *rank projection tree* method with different prioritization functions to pick out individual genes relevant to the diseases predicted by each network. For this purpose, we use a simple ranking function where, for layers *l* and *m* = *l* + 1, *r*_*i*_,_*l*_,_*m*_(*j*) returns the rank of *n*_*m*_,_*j*_ in the ordering induced by the weights *W*_*l*_,_*m*_(*i*,.) (signed values, descending), and set *B* = 2. For the prioritization function *π*, we first calculate the cumulative rank-scores *c*_*t*_ of each path in *T*; that is, for leaf node *t* = [*b*_1_,…*b*_*L*_], we set *c*_*t* = (∑_*l*_|*b*_*l*_|) (∏_*l*_ sign(*b*_*l*_)). We then set 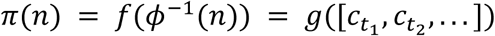 for *t*_1_,*t*_2_… ∈ *ϕ*^−1^(*n*), where *g*(.) is one of the functions: {sum, average, max, min} of signed or absolute values, or length. Table 1 scores the rankings of the top 20 genes (averaged across networks) according to each *g*(.) when compared against a ‘ground truth ranking’ based on the number of citations retrieved by Google Scholar when each gene is queried in association with the disease; the table shows the *ℓ*_1_-distance between the two rankings, normalized by its maximum (*ℓ*_1_([1…20], [20…1]_*m*_). In general, the absolute average and max functions appear to be better indicators of individual gene importance.

**Table 1.**
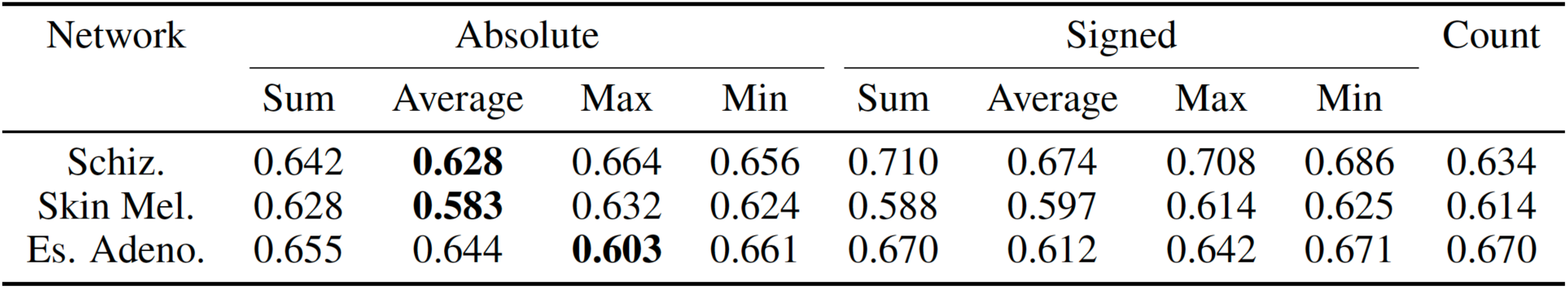
Prioritization function comparison. The rankings of genes induced by different prioritization functions are compared against citation-based rankings from existing literature. Table shows normalized *ℓ*_1_-distances of predicted and citation-based rankings for the top 20 genes, with best performing metrics highlighted. Rows are: Schizophrenia, Skin Melanoma and Esophageal Adenocarcinoma.

| Network | Absolute |  |  |  | Signed |  |  |  | Count |
| --- | --- | --- | --- | --- | --- | --- | --- | --- | --- |
|  | Sum | Average | Max | Min | Sum | Average | Max | Min |  |
| Schiz. | 0.642 | **0.628** | 0.664 | 0.656 | 0.710 | 0.674 | 0.708 | 0.686 | 0.634 |
| Skin Mel. | 0.628 | **0.583** | 0.632 | 0.624 | 0.588 | 0.597 | 0.614 | 0.625 | 0.614 |
| Es. Adeno. | 0.655 | 0.644 | **0.603** | 0.661 | 0.670 | 0.612 | 0.642 | 0.671 | 0.670 |

### 3.2 Multigene groupings

To test the ability of the *rank projection tree* to extract meaningful gene groupings at multiple levels, we extracted all positive and negative groupings across layers *l* = 0…3 using Eq. 2 for the schizophrenia networks, using exactly the same settings as above. For comparison, we built identical trees, but used a randomized ranking function *r*(.) to produce the *ϕ* mapping to the schizophrenia networks. We used the networks with the WGCNA modules average expression levels as inputs; hence, for set *S* formed from Eq. 2, we take the union of the genes in all modules which are elements of *S*.

For our first comparison, we annotate all groupings with KEGG pathway terms using gene-set enrichment analysis [32]. All terms are assigned to a grouping achieving a q-value *<* 0.001, and a ranking across KEGG terms is made for each layer independently by counting the number of groupings a term is associated with across all models (including duplicate groupings, hence accounting for increased importance of nodes in *N* mapped to by multiple nodes in *T*). We take the top 20 such terms, and sum the number of citations associated with these terms in association with schizophrenia by Google Scholar as above. Table 2A compares the total citations across layers, showing that the rank projection tree finds more important terms than the random tree at all layers, and the groupings produced by higher layers of the network (*L*_3_ ‘lowest’, *L*_0_ ‘highest’) tend to associate with more trait-relevant terms (peaking at the penultimate layer, *L*_1_).

**Table 2.** Multigene and subject groupings in psychiatric genomics data. (A) Top compares the total citations of the top 20 KEGG terms associated with *rank projection tree* and randomized gene groupings at each layer of neural networks trained to predict healthy versus schizophrenia with two hidden layers. (B) Bottom compares the number of subject clusters discovered by *cascaded network decomposition* found to be associated with each semantic category. Associations were determined by a 0.1 threshold on the *p*-value of a hypergeometric test, and the totals are counted across models trained on 10 data-splits.

| Model | $L_0$ | $L_1$ | $L_2$ | $L_3$ |
| --- | --- | --- | --- | --- |
| Rank-projection | 65K | 82K | 77K | 73K |
| Randomized | 53K | 54K | 56K | 72K |

**Table 2. Multigene and subject groupings in psychiatric genomics data.**
| Category | SCZ | BPD | ASD |
| --- | --- | --- | --- |
| Gender | 4 | 2 | 1 |
| Age | 7 | 1 | 2 |
| Ethnicity | 2 | 1 | - |
| Psychotic Med. | - | 1 | - |
| ASD Subtype | - | - | 2 |

Finally, we analyzed the enrichment of the groupings from all layers as described above for ‘high-confidence schizophrenia genes’. These are genes which can be linked to GWAS hits for schizophrenia by any three of the following four methods: Hi-C interactions; enhancer-target links (using covariation of chromatin and expression signals); eQTL linkages, and isoform-QTL linkages (321 genes; list to be made available in [2]). The enrichment of such genes in each module is scored using a p-value from the hyper-geometric test. Fig. 4A shows that these genes are significantly more enriched in the groupings found by the rank projection tree than randomized trees, where the groupings found by the penultimate layer *L*_1_ again appear most trait-relevant, matching the findings of the citation-based metric above. We also compared the *L*_3_ distribution with a gradient-based prioritization scheme which ranked the modules at this level according to the absolute magnitude of the gradient of the network output with respect to each input (as in [7]), but found that it was not significantly better than the randomized tree (p=0.78); the rank projection tree was better than the randomized tree across all layers (p<1e-4 for *L*_1−3_, and p=0.012 for *L*_0_, all p-values using 1-tail KS-tests).

**Figure 4.**
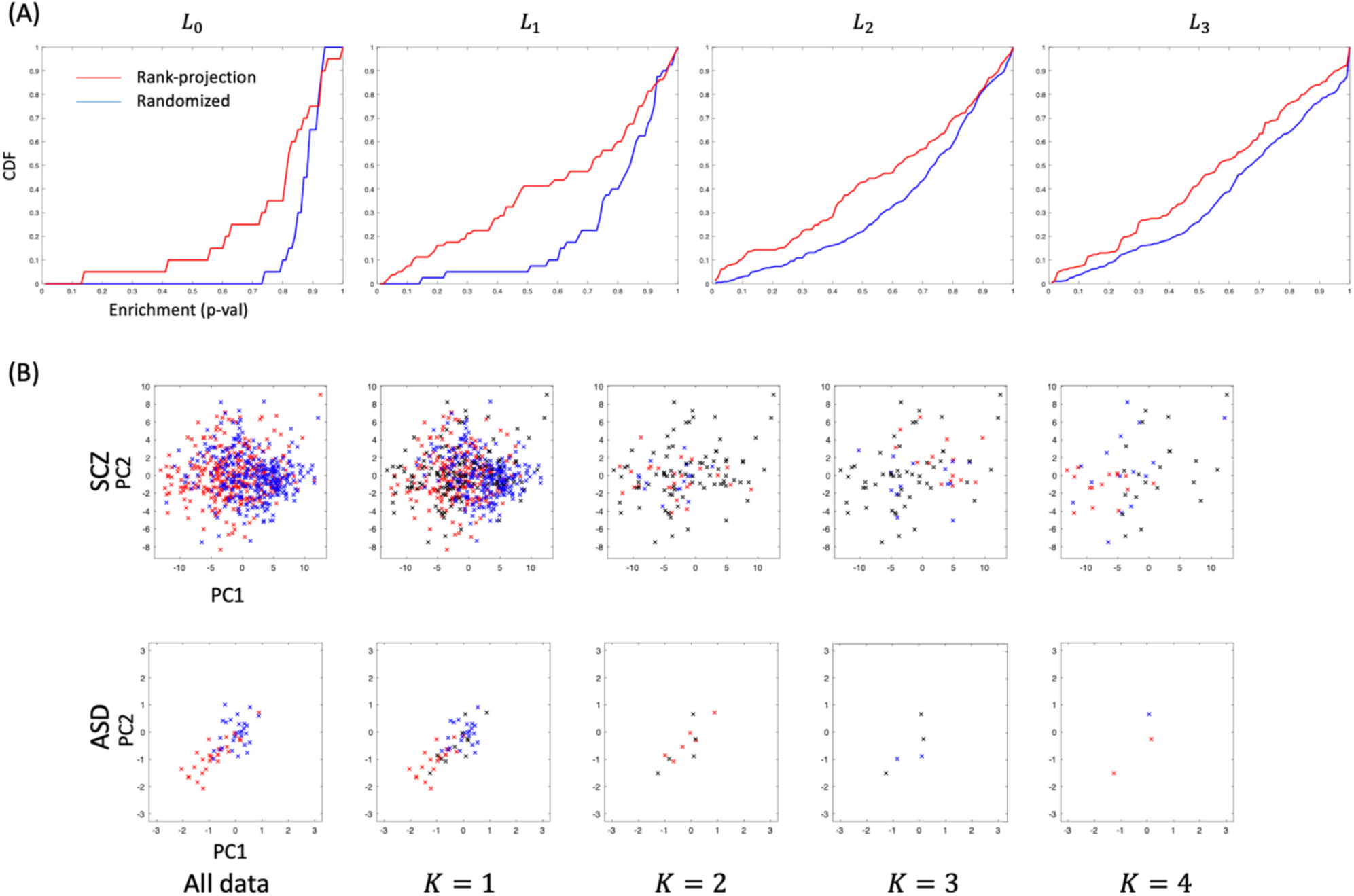
Visualization of network interpretations. (A) enrichment of high-confidence schizophrenia associated genes in gene groupings found associated with different neural network layers. Enrichment p-values are from the hyper-geometric test, and the empirical cumulative density function (CDF) is plotted on the y-axis. (B) compares the subject groupings derived from different levels of the cascaded network decomposition applied to SCZ and ASD subjects. Subjects are projected into PCA-space using their gene expression profiles; red, blue and black denote cases, controls and rejected points at each stage of the cascade (mirroring Fig. 3B).

### 3.2 Instance-based groupings

We next test the ability of the *cascaded network decomposition* approach to extract meaningful instance/subject groupings from the PsychENCODE psychiatric genomics dataset, using the same data and network settings as above for Schizophrenia (SCZ), in addition to 10 data splits each for Bipolar (BDP) vs control subjects, and Autism Spectrum Disorder (ASD) vs control subjects (see Appendix). We compress the networks trained on each data split, and extract subject groupings for each stage of the cascade in the compressed networks. We analyze the extracted subject groupings for enrichment of semantic categories, including Gender, Age (binarized at the median), Ethnicity, as well as treatment with Psychotic medication and ASD subtype (see [2]).

Fig. 4B visualizes the subject groupings extracted from two cascades trained on SCZ and ASD datasets respectively. Notably, in both cascades, while the case-control axis for the groupings associated with the first stage of the cascade (*K* = 1) is the same as the dominant axis for the dataset, in the second stage (*K* = 2) the axis is reversed, suggesting that these subjects are characterized by different criteria. Table 2B then provides a breakdown of the number of groupings enriched for each semantic category across all datasets. Notably, the SCZ groupings are more highly enriched for semantic groupings, which may reflect the increased size of the SCZ dataset. Further, we note that signal for enrichment in subjects prescribed psychotic medication is weaker than the other categories, in agreement with prior observations that the effects of medication on the post-mortem transcriptome are weak in this dataset [31]. Finally, the ability of the cascade approach to find groupings associated with ASD subtypes reflects the associations of these subtypes with PFC and other brain regions [31]; the models are trained on PFC expression data, leading to an association of the PFC subtype with earlier stages in the cascade, and the non-PFC subtype with later stages (see Appendix A).

### 3.4 Compression and Generalization

Secs. 3.1-3 have shown that the RPT and CND methods are able to extract semantically meaningful gene and subject groupings from the datasets tested. For this purpose, we took a global view, exploring the characteristics of the groupings extracted across models, data-splits and cascade stages, while fixing the compression branching factor in the case of the RPT. Here, we consider the interpretations extracted at a finer-grained level; particularly, we investigate whether there is a relationship between the strength of the semantic associations and the generalization of the network, and if the compression bounds in Sec. 2.3 can be used to predict such a relationship (hence we test whether models which generalize better, or are predicted to do so, are more interpretable).

We begin by investigating the strength of association between the PAC-Bayes bound (Eq. 8) with the MDL-prior (Eq. 9) in predicting the test error for both the RPT and CND compression schemes. For this purpose, we use RPTs to derive compressed networks of varying degrees of compression by setting the half-branch factor to *B* = {1,2, … 5} for networks trained on each of the 10 SCZ data splits. We evaluate the bound in Eq. 8 by compressing the parameters of the resulting networks using LZW compression [33], and use the resulting binary code as the network representation 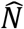 in Eq. 9. Similarly, we use the cascades learned by CND on each of the psychiatric datasets to derive a series of compressed models, by truncating each cascade after each possible stage, and compressing the resulting models by LZW compression to generate the binary codes, 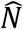. Fig. 5A and 5B show plots of the correlation between the empirical test error and the MDL bounds calculated as above using the RPT and CND schemes respectively. In Fig. 5A, the bound parameter *λ* is fitted independently for each group of compressed models associated with a given uncompressed model (corresponding to a fixed data split), hence each plot contains networks associated with *B* = {1,2, … 5}. In Fig. 5B, similarly, *λ* is fitted independently for the truncated models associated with each cascade, hence the number of datapoints is *K*, the length of the full cascade. We further standardize each bound, by dividing by the variance per plot, and subtracting the minimum distance with the test error across all models; these steps represent an ‘empirical normalization’ of the bound, and are for visualization purposes only, since we are only concerned with correlations with the test error. A sign-test on the correlations between bounds and test error shows that there is significant positive correlation (*p* = 0.022 and *p* = 0.031 for RPT and CND schemes respectively). Further, since we fit *λ* independently on each group of models, we additionally test the informativeness of the MDL component of the bound, by comparing against a null distribution, in which we permute the model description lengths (the binary codes) and optimize *λ* as above for 20 permutations per model. Using the mean correlation over the top 3 models as the test statistic, this gives *r* = 0.954 and *r* = 0.926 for RPT and CND schemes respectively, which are significant at the *p* = 0.05 and *p* = 0.2 levels according to the permutation test. We then investigate the relationship between generalization and interpretability for each compression scheme. For the RPT scheme, we directly correlate the predicted accuracy (defined as 1 − MDL-bound) for each model with the KS test statistic, calculated as in Sec. 3.2 for the gene groupings derived per-layer from the RPT compressed networks (based on the enrichment of high-confidence SCZ genes from [2]). Fig. 5C shows the correlations across all models, which are significantly skewed positive (*p* = 0.0072, 1-sample t-test). Further, we observe from Sec. 3.2 that the degree of positive correlation is strongest in layer L0, and decreases for subsequent layers, mirroring the variation across layers shown in Fig 3.A. For the CND scheme, we divide the subject groupings into ‘semantically enriched’ and ‘non-enriched’ categories, corresponding to those associated with the categories in Table 2B. For both these groups, we compare the change in test-error, or MDL-bound, which results from adding the cascade stage from which the grouping derives to the model. The rationale for this, is that the change in predictive power may serve as an approximate measure of the ‘informativeness’ of the local boundary found by a given cascade stage, and hence the semantic relevance of the groupings associated with the stage in question. Fig. 2D shows that there is indeed an enrichment in larger reductions in test-error associated with semantically relevant groupings; while the trend is in the same direction for the MDL-bound, the effect is not significant, suggesting that a tighter bound may achieve a stronger correspondence (for instance, using a larger dataset).

**Figure 5.**
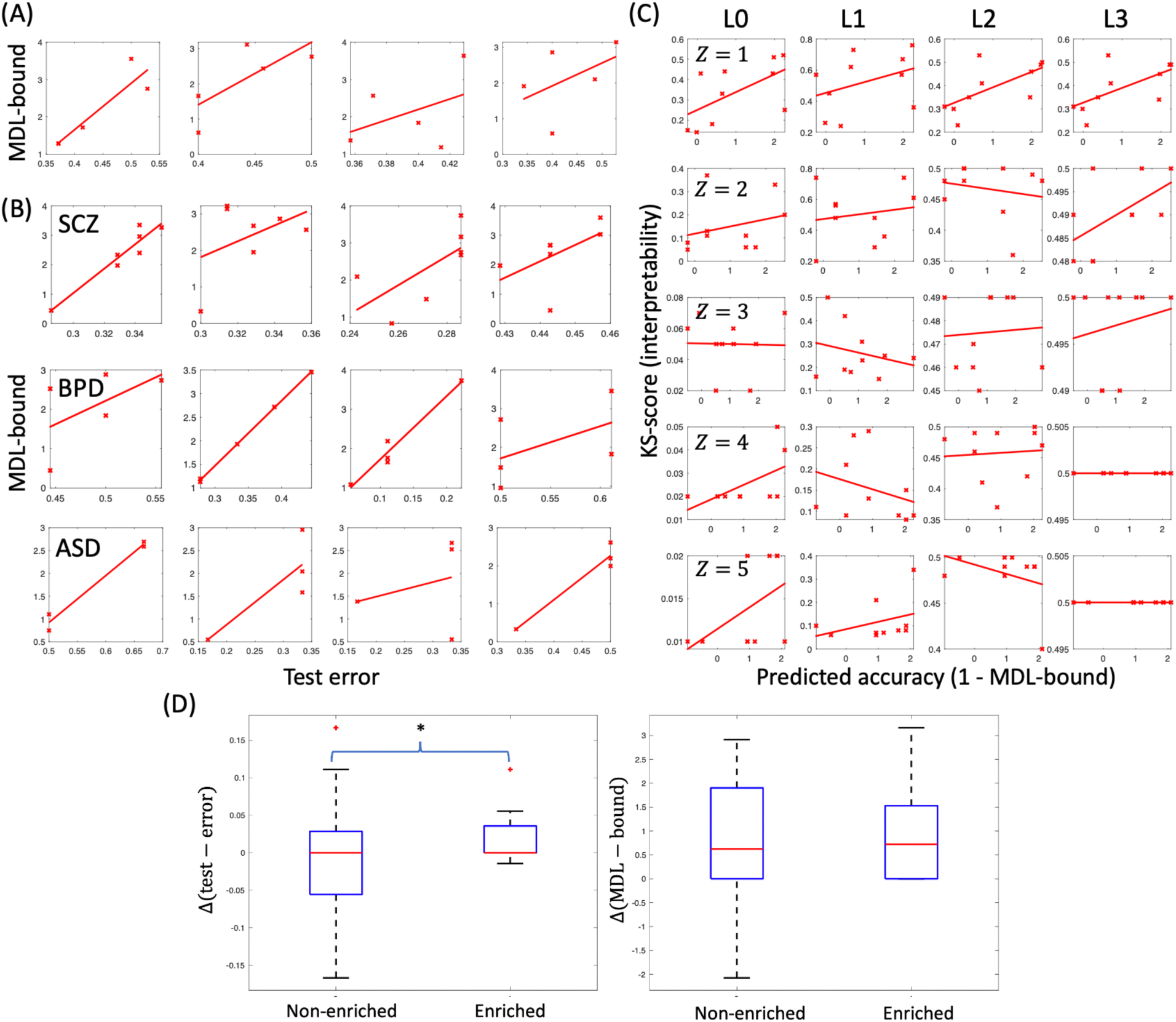
Predicting semantic enrichment using generalization bounds. (A) and (B) plot test error vs. MDL bounds for groups of models derived from the RPT and CND schemes respectively. (C) shows the relationship between interpretability and predicted accuracy (defined as 1 − MDL-bound) per network layer (*L*) and compression strength (*Z*) for the RPT. (D) shows the relationship between per-cascade-stage generalization and semantic enrichment for the CND scheme. See text for details.

Finally, we investigate the modified MDL bound introduced in Eq. 10, combining both MDL and data-dependent components, using the RPT scheme. We replicate the generalization test from Fig. 5A using this bound, while fitting *λ* and *α* for each group of models. This achieves an improved mean correlation of *r* = 0.955 for the top 3 models, which is again significant at the *p* = 0.05 level (permutation test). Table 3 further compares the mean KS-bound correlation across models (as in Fig. 5C) of the MDL and modified-MDL bound, showing only a marginal increase in correlation. Table 3 further shows that both bound achieve a significantly stronger correlation with the semantic KS-scores than the observed test error; a possible reason for this is the use of the training error in the first term of the bound, which is typically more stable than the test error, given the larger number of data points (640 vs. 70 in the SCZ datasets).

**Table 3.** Comparing MDL and Modified MDL bounds for predicting semantic enrichment. Table shows mean Pearson correlation and 1-sample t-test p-values for models using the RPT scheme, when the KS-semantic-enrichment is correlated with the quantities shown.

| Bound | Test error | MDL | MDL+prior |
| --- | --- | --- | --- |
| mean $r$ | 0.142 | 0.217 | 0.218 |
| $p$-val | 0.025 | 0.007 | 0.007 |

## 4. Discussion

We have introduced the general framework of a *network interpretability scheme*, based on model transformation (particularly compression), information extraction, and an ‘interpretable model selection’ step. Further, we have introduced two complementary schemes using model compression, rank projection trees, and cascaded network decomposition, which allow feature groups and data instance groups to be extracted from a trained network that may have semantic significance. We further outline a general purpose MDL generalization bound suitable for such compression schemes (based on [15]), which we extend to a modified MDL bound with MDL and data-dependent components. We show that both generalization performance and the bounds we introduce are predictive of semantic enrichment, with a small benefit afforded by the modified bound, strengthening our claim that compression an generalization analysis should be combined in order to derived optimal interpretations of a network (one which optimally preserves the network’s *implicit semantics* [16]).

A number of future directions are suggested by our work. One possibility is to extend our consideration of combined MDL and data-dependent bounds to explicitly handle cases where prior structure is built into network models. For instance, in [2], the structure of a gene regulatory network is explicitly encoded into a deep network (a Deep Structured Phenotype Network, DSPN), and novel gene groupings are extracted by a compression scheme similar to our RPT. A formal analysis of generalization in such cases may be achieved by including structural features of the network in the data-dependent prior, which in turn may motivate enhanced interpretation methods using by selecting groupings based on the corresponding bound.

We also note that, while we have concentrated on schemes which do not explicitly constrain the resulting model class (beyond sparsity), such constraints may be straightforwardly embedded as in [13], and a targeted form of compression may be achieved by optimizing for generalization within a restricted model class whose structure may be readily interpreted in an externally defined context (for instance, decision trees as a basis for decision making).

Finally, while we have concentrated on genomics applications, the compression schemes we introduce, as well as the associated bounds may be readily investigated in other domains such as image procession and language. We expect a similar relationship to hold between interpretability and MDL-based generalization schemes in other domains, although the optimal bounds for performing interpretable model selection may vary greatly across domains, for instance in response to different kinds of relevant prior information.

## A. Appendix A

### A.1 Proof of Theorem 1

#### Theorem 1 (Modified MDL Bound).

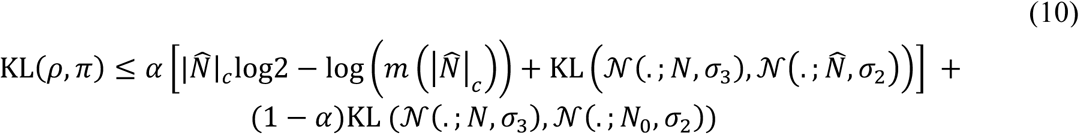

*Proof*. We can decompose the LHS into cross-entropy and Shannon entropy terms:

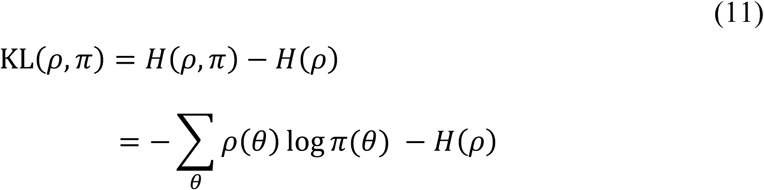

Using Jensen’s inequality, we have:

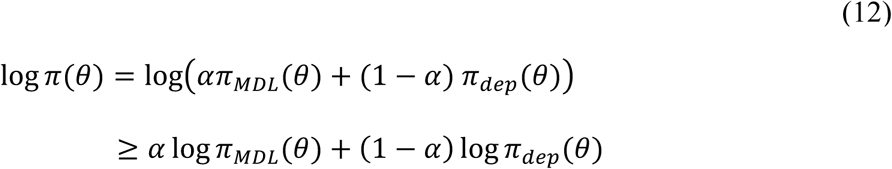

Hence, substituting into Eq. 11:

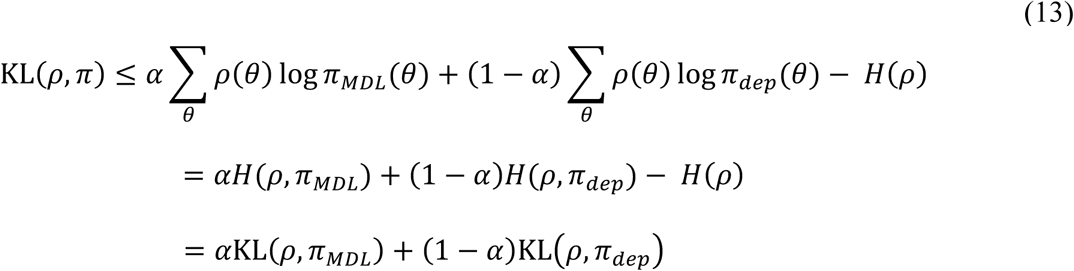

The theorem follows by substituting the result of Th. 4.3 from [15] into the first term of Eq. 13.□

### A.2 Datasets and Network details

#### PCAWG

The PanCancer Analysis of Whole Genomes (PCAWG) study includes a variety of biological data types corresponding to 2,800 samples from the International Cancer Genome Consortium. To train networks for our analysis, rare variants are singled out for Skin Melanoma and Esophageal Adenocarcinoma samples. The predictive task according to which the neural networks have been trained is the prediction of somatic and germline variation co-occurrence at the gene level for 718 genes of the COSMIC census list fetched on May 08, 2018. Input data included 43 features ranging from germline variant signatures of known cancer genes alongside a set of biological features extracted from multiple data and annotation repositories, namely UCSC Genome Browser [34], Gencode v27 [35], and COSMIC [36]. Each model whose weights have been analyzed by *rank projection trees* has 3 hidden layers. Number of hidden nodes (285-941), optimization algorithm (Adam or Nesterov Adam), and activation functions (Exponential or Rectified Linear Unit) for each network have been determined by automated hyperparamter optimization using the HyperOpt package [37]. Results are averaged for 5 neural networks trained on randomly stratified training datasets for each cancer type, with test performance of high precision and recall values ranging between 70% and 83%. To balance training datasets, we deployed the SMOTE oversampling algorithm [38] using the implementation in the imbalanced-learn Python package [39].

#### PsychENCODE

The PsychENCODE dataset [30,2] contains bulk transcriptomics and other omics data from the prefrontal cortex of 1452 post-mortem subjects, including controls and subjects with schizophrenia, bipolar disorder, and autism. From these data, we create 10 training and testing partitions (including 640 and 70 samples respectively) of control and schizophrenia subjects, which are balanced 50-50% for controls and cases, and balanced across train/test partitions for covariates including age, gender, ethnicity and assay. We train neural networks with 2 hidden layers to predict a binary case/control indicator, with 100 and 400 nodes at layers 1 and 2 respectively, logistic sigmoid activations, and SGD with early stopping for training. We train separate neural networks using individual gene expression levels as inputs, and mean expression levels across modules of genes pretrained using WGCNA [22,23], pre-selecting the top 1% and 15% of genes/modules respectively according to the absolute Pearson correlation between the input and the binary output indicator on each training partition (resulting in 187 genes and 754 modules in each respective model). The test performance of the models averaged across partitions was 73.6% and 66.1% for the gene- and module-based models, respectively.

